# Telocytes link epithelial nutrient sensing with amplification of the ILC2-tuft cell circuit

**DOI:** 10.1101/2024.10.14.618111

**Authors:** Chang Liao, Ming Ji, Zhi-En Wang, Daniel J. Drucker, Hong-Erh Liang, Richard M. Locksley

## Abstract

Group 2 innate lymphocytes (ILC2s) are prevalent in small intestine but engagement of type 2 immunity during basal processes are incompletely described. Thymic stromal lymphopoietin (TSLP), a cytokine implicated in ILC2 activation, was constitutively expressed in villus telocytes and crypt-associated trophocytes, specialized fibroblasts that sustain epithelial identity. Feeding increased TSLP and induced ILC2 type 2 cytokines that were attenuated by deletion of TSLP in PDGFRα^+^ stromal cells or TSLP receptor on ILC2s. Mouse and human telocytes expressed receptors for glucagon-like peptide-2 (GLP-2), which is released by enteroendocrine cells (EECs) after eating. GLP-2 induced intestinal TSLP, TSLP-dependent ILC2 cytokine production, and tuft cell hyperplasia. The telocyte-alarmin relay couples EEC nutrient detection with amplification of a tuft cell chemosensory circuit that diversifies surveillance of ingested cargo.

**One-Sentence Summary:** Intestinal telocyte TSLP relays signals from enteroendocrine cells to ILC2s to amplify the tuft cell circuit in response to feeding.

## Main Text

The small intestinal mucosa represents a key barrier balancing host nutritional needs with exposure to potentially noxious chemical and infectious environmental agents. Innate and innate-like lymphocytes, including group 3 innate lymphoid cells (ILC3s), intraepithelial lymphocytes (IELs), γδ T cells (*1–4*) and specialized RORγt^+^ antigen-presenting cells (*5*), create a healthy IL-17/22-mediated and regulatory T cell (Treg)-controlled type 3 immune tissue environment that sustains the dynamic interface facilitating nutrient absorption, metabolism, microbiota control, tolerance and epithelial protection during fasting, feeding and circadian cycles. Innate type 2 immune cells, including ILC2s, are abundant in small intestinal lamina propria (siLP), and, with adaptive Th2 cells, are highly engaged during responses to intestinal helminths and in food allergy. Immune responses to intestinal helminths are only partially effective resulting in high prevalence of infection and frequent recurrences after therapy in exposed populations. Food allergy represents a mis-directed immune attack against non-harmful ingested constituents. Neither of these conditions reveals an underlying pathway mediated by resident type 2 immune cells in small intestine that becomes overtly prominent during parasitic and allergic pathology.

In prior studies, our laboratory and others identified a key role for the alarmin IL-25 as a tuft cell-derived cytokine that activates siLP ILC2s to release IL-13, which biases epithelial progenitors to a secretory cell fate that increases goblet and tuft cells in a forward-amplifying circuit accounting for the ‘weep and sweep’ response to helminths and protists (*6–8*). Deletion of cell intrinsic regulators of IL-25 signaling, such as *Tnfaip3* (A20), resulted in constitutive activation of the circuit, suggesting control of the basal state (*9*). The combinatorial role of alarmins in regulating the activation of tissue resident ILC2s and Th2 cells (*10,11*) prompted us to consider roles for additional alarmins in small intestinal physiology. Here, we demonstrate that thymic stromal lymphopoietin (TSLP) from specialized subepithelial stromal cells relays signals from epithelial enteroendocrine cells (EECs) to siLP ILC2s to amplify the tuft cell circuit by a reversible process responsive to feeding and fasting.

### Specialized subepithelial fibroblasts express TSLP in small intestine

We targeted the *Tslp* locus to identify TSLP-producing cells using the flox-and-reporter (Flare) cassette used previously to visualize IL-25-producing tuft cells (*6*) (Fig. 1A). Under basal conditions in specific pathogen-free C57BL/6 Flare-TSLP mice, the predominant TSLP reporter-positive (TSLP-tdTomato^+^) cells in small intestine were subepithelial PDGFRɑ^+^Podoplanin (PDPN)^+^ cells with the morphology of fibroblasts enriched at the villus tips throughout small intestine and organized within crypt-associated cellular networks in the jejunum and ileum (Fig. 1B, fig. S1A, Movie S1-S7). These cells comprised >80% of TSLP-tdTomato^+^ cells among non-hematopoietic cells and increased in total prevalence from proximal (1-4%) to distal small intestine (2-7%). TSLP-tdTomato^+^CD31^+^PDPN^+^ lymphatic endothelial cells were fewer, comprising 10-20% and 1-10% of TSLP-tdTomato^+^CD45^-^ cells in proximal and distal small intestine, respectively. TSLP-tdTomato^+^CD45^+^ cells were found only among rare CD31^+^ endothelial cells (Fig. 1C-D, fig. S1B-D). Despite reports of TSLP-positive small intestinal epithelia (*12,13*), including type 2 tuft cells, no intestinal TSLP-tdTomato^+^ epithelial cells were identified by flow cytometry or microscopy in Flare-TSLP mice (fig. S1F and Fig.1B). This was not due to an inability to visualize epithelial cell TSLP since topical application of the vitamin D analog calcipotriol (MC903), a known inducer of TSLP in keratinocytes (*14*), induced reporter signal in Flare-TSLP mice (fig. S1E). In small intestine, epithelial tuft cells remained TSLP reporter-negative during a kinetic analysis of acute *Nippostrongylus brasiliensis* infection (fig. S1F-G).

**Fig. 1.**
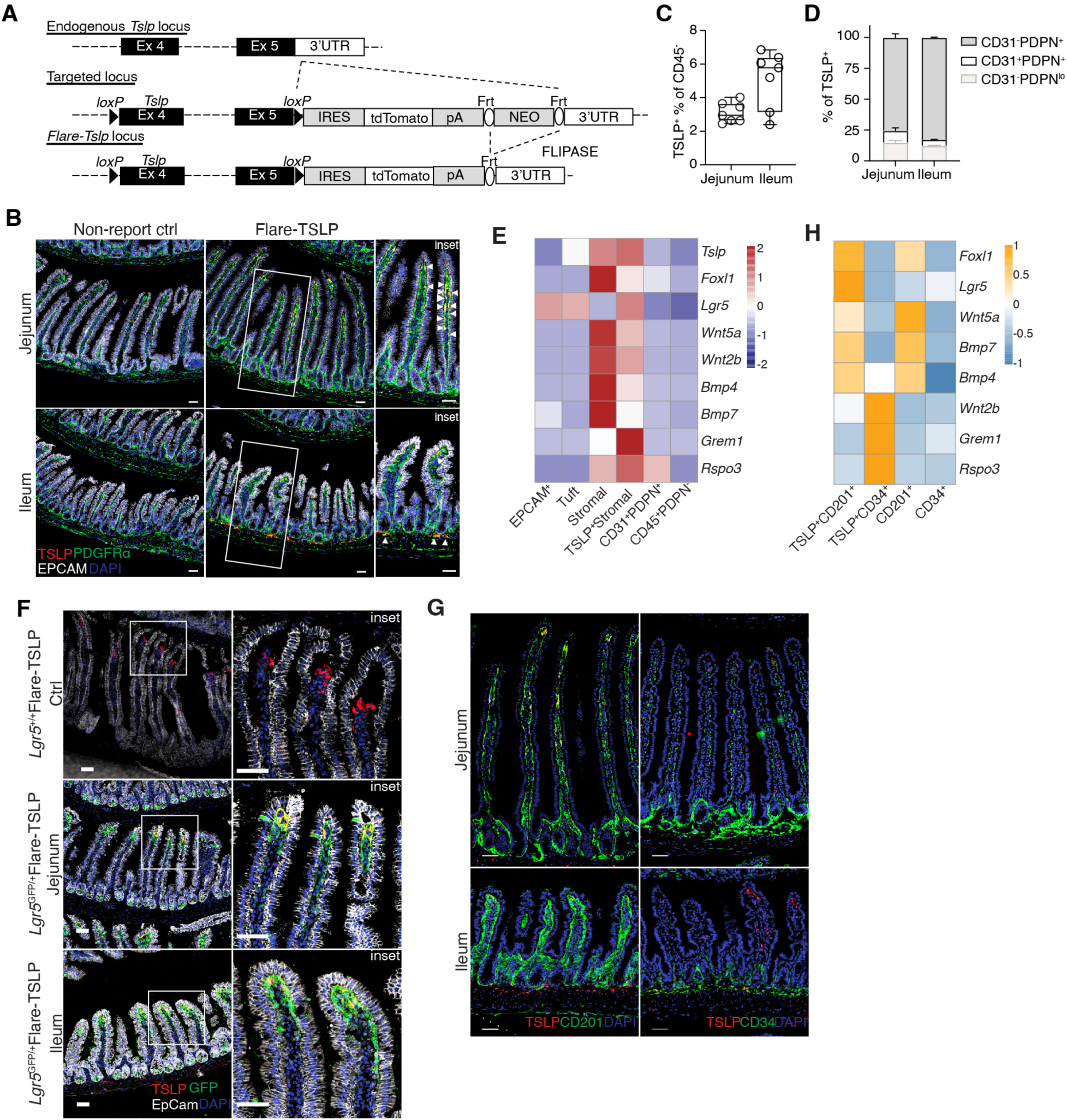
Telocytes and trophocytes are primary sources of TSLP in the small intestine. **(A)** Gene-targeting strategy for the flox and reporter of *Tslp* (Flare-TSLP) mouse. Frt, target site for FLIPASE recombinase; IRES, internal ribosomal entry site; *loxP*, target site for Cre recombinase; pA, bovine growth hormone poly(A) tail; tdRFP, tandem-dimer red fluorescent protein, or tdTomato; UTR, untranslated region. **(B)** Representative imaging of jejunum (**top**) and ileum (**bottom**) of Flare-TSLP mice or non-reporter control mice. Red, TSLP-tdTomato; Green, PDGFRɑ; White, EPCAM; Blue, DAPI. 20x objective. Arrowheads indicate TSLP-tdTomato^+^ cells. Scale bar, 50 µm. (**C**)(**D**) Flow cytometric analysis of endothelial and other stromal populations in jejunum and ileum small intestine lamina propria (siLP) in Flare-TSLP mice. (**C**) Percentage of total TSLP-tdTomato^+^ in CD45^-^ population. (**D**) Percentage of endothelial cells and other stromal within TSLP-tdTomato^+^ cells. Cells were stained with anti-CD31 and anti-Podoplanin (PDPN) antibodies. Lymphatic endothelial cells are CD45^-^PDPN^+^CD31^+^ and fibroblasts are CD45^-^ PDPN^+^CD31^-^. **(E)** RT-qPCR analysis of purified CD45^-^EPCAM^+^cells, CD45^-^EPCAM^+^ SiglecF^+^ tuft cells, CD45^-^CD31^-^PDPN^+^ total stromal cells, CD45^-^CD31^-^PDPN^+^TSLP-tdTomato^+^ cells, CD45^-^CD31^+^ PDPN^+^ lymphatic endothelial and CD45^+^PDPN^-^ hematopoietic cells from small intestinal epithelial fraction or siLP (whole tissue) of Flare-TSLP mice. Gene expression normalized to *18S*. **(F)** Representative imaging of jejunum (**middle**) and ileum (**bottom**) of Flare-TSLP; *Lgr5*-eGFP dual reporter mice or Flare-TSLP single reporter (Ctrl) mice (**top**). Red, TSLP-tdTomato; Green, *Lgr5*-eGFP; White, EPCAM; Blue, DAPI. 20x objective. Scale bar, 50 µm. **(G)** Representative imaging of CD201 and CD34 expression in jejunum (**top**) and ileum (**bottom**) of Flare-TSLP mice. Red, TSLP-tdTomato; Green, CD201 (**left**) or CD34 (**right**); Blue, DAPI. 20x objective. Scale bar, 50 µm. **(H)** RT-qPCR analysis of purified TSLP-tdTomato^+^CD201^+^CD31^-^ (telocytes), and TSLP-tdTomato^+^CD34^+^CD31^-^ (trophocytes), CD201^+^CD31^-^, and CD34^+^CD31^-^ stromal cells from siLP (whole tissue) of Flare-TSLP mice. Gene expression normalized to *18S*. All data are biological replicates, n≥3. Data are representative of at least two independent experiments. Error bars indicate samples mean± SEM.

The location and morphology of TSLP^+^ fibroblasts in Flare-TSLP mice resembled previous reports of subepithelial telocytes and crypt-associated trophocytes, which are structural cells that generate the counter-regulating WNT and BMP gradients that maintain the crypt stem cell niche and the axis controlling epithelial differentiation during vertical zonation of the villi (*15,16*). To confirm the identity of these cells, we used fluorescence-activated cell sorting (FACS) to purify TSLP-tdTomato^+^ cells and used RT-qPCR to assess expression of signature transcripts. CD45^-^CD31^-^PDPN^+^TSLP^+^ stromal cells expressed peri-villus telocyte markers including *Foxl1*, *Wnt5a* and *Bmp7*, as well as crypt-associated trophocyte markers including *Wnt2b*, *Grem1* and *Rspo3* (*15*, *16*), whereas the expression levels of these genes were relatively low in EPCAM^+^ total epithelial cells, tuft cells, CD45^-^CD31^+^PDPN^+^endothelial cells or CD45^+^ PDPN^-^ hematopoietic cells, except for *Rspo3*, which was also expressed by endothelial cells (Fig. 1E). Expression of *Lgr5* in the TSLP^+^ population (Fig.1E) suggested that villus tip telocytes (VTTs), specialized telocytes that serve as signaling hubs for specifying epithelial cell fate, express TSLP (*17*). Using Flare-TSLP; *Lgr5*-eGFP dual reporter mice we confirmed co-expression of *Tslp* and *Lgr5* among villus telocytes, consistent with their identification as VTTs (Fig. 1F). Prior studies identified CD201 and CD34 as markers for intestinal telocytes and trophocytes (*16,18*), respectively. In line with this, CD201 labeled villus and *Lgr5*-GFP^+^ telocytes but not trophocytes, whereas CD34 labeled crypt-associated TSLP^+^ trophocytes (Fig. 1G and fig. S2A-C). We purified CD31^-^CD201^+^TSLP^+^ and CD31^-^CD34^+^TSLP^+^ cells (fig. S2D) and confirmed their expression of signature transcripts of telocytes and trophocytes (Fig. 1H and fig. S2E). Thus, specialized fibroblasts designated telocytes, including VTTs, and trophocytes constitute the primary cell types that express TSLP in mouse small intestine under resting conditions.

### Small intestine TSLP increases with feeding and activates ILC2 cytokine production

Feeding activates type 2 cytokine expression by small intestine ILC2s independent of the light-dark cycle, in part through VIP-mediated IL-5 production in control of eosinophil homeostasis (*19*). After 16 hr fasting, Flare-TSLP mice were gavaged orally with a 500 μl slurry of powdered diet containing ∼2 Kcal mixed nutrients (∼1% total daily caloric intake with nutritional components in proportion to standard chow diet) or water control (Fig. 2A). After 2 hr, the prevalence of TSLP-tdTomato^+^ CD45^-^ nonepithelial cells increased in proximal and distal siLP (Fig. 2B and fig. S3A). We confirmed TSLP protein upregulation by ELISA using jejunal and ileal tissue explants and homogenates. TSLP increased and peaked around 2 hr after gavage (fig. S3B). TSLP upregulation at 2 hr after feeding was unchanged in fasted *Pou2f3*^-/-^ mice that lack tuft cells, suggesting that these cells were not required (Fig. 2C and fig. S3C). To assess the role for TSLP in ILC2 activation after feeding, we used cytokine reporter mice (*20*) containing marker alleles for IL-5 (Red5) and IL-13 (Smart13) crossed to TSLPR-deficient (*Tslpr*^-/-^) mice and analyzed ILC2s from proximal jejunum after 16 hr fasting and 4 hr refeeding with chow diet and water *ad libitum* or water alone (Fig. 2D). In line with prior findings (*19*), ILC2s were activated 4 hr after feeding as assessed by an increase in KLRG1^+^IL-5(Red5)^+^IL-13 (Smart13)^+^ cells that was diminished in TSLPR-deficient mice with almost complete loss of IL-13 reporter expression (Fig. 2E-F). We next generated *Tslp^fl/fl^Pdgfrɑ^CreERT2^*Smart13 reporter mice. Following injection of tamoxifen, these mice delete TSLP in PDGFRɑ-expressing stromal cells, allowing us to assess contributions of stromal TSLP to activation of intestinal ILC2s by food intake. Loss of TSLP from stromal cells caused attenuation of ILC2 activation as assessed by diminished expression of the IL-13 reporter after feeding (Fig. 2G-H). Loss of ILC2 activation after feeding was also observed in *Tslpr^fl/fl^Il5^Cre+^*Smart13 mice with deletion of TSLPR on IL-5-expressing intestinal ILC2s (Fig. 2I), demonstrating that stromal cell-derived TSLP was needed to activate ILC2s through TSLPR after feeding.

**Fig. 2.**
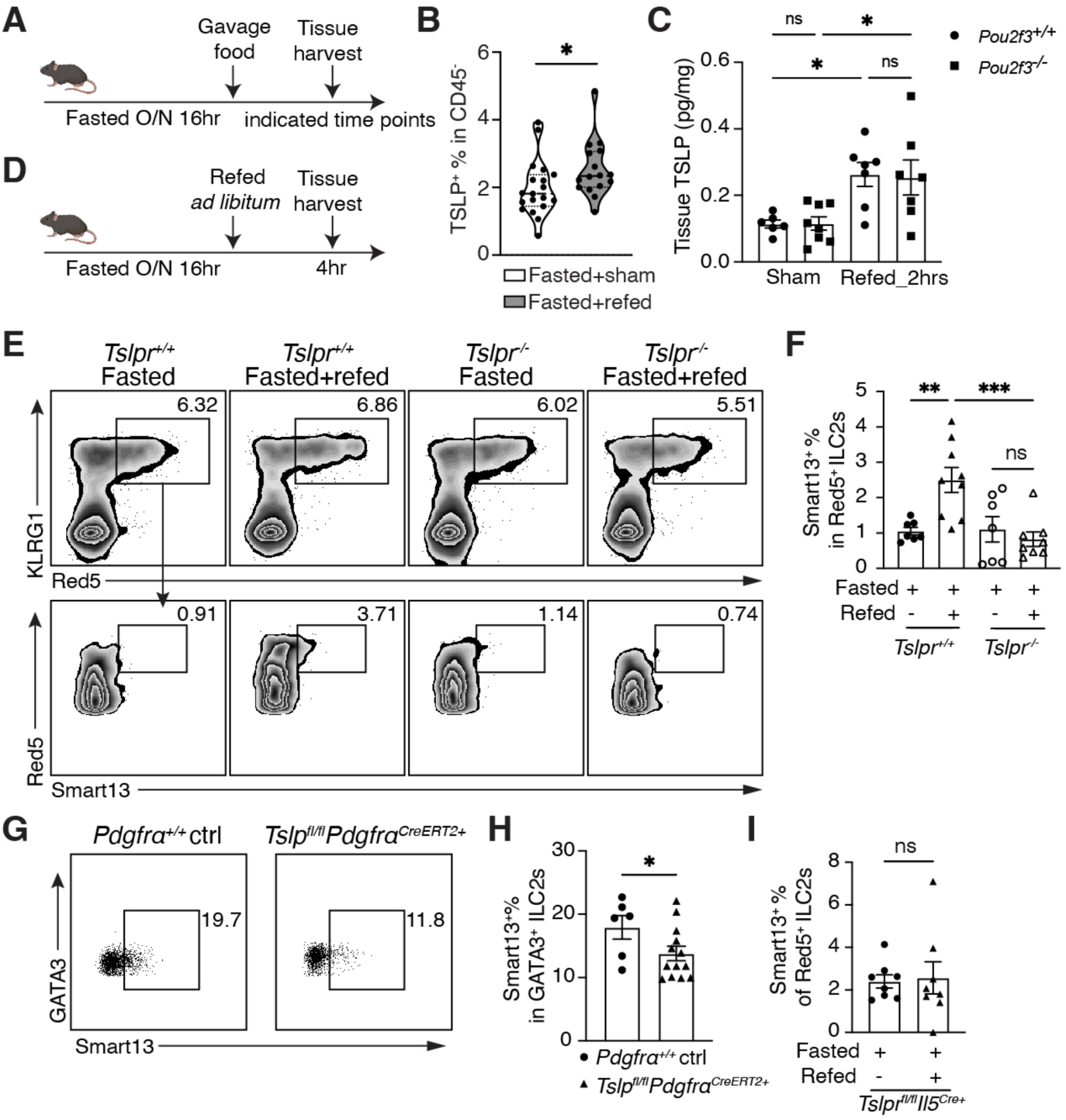
Feeding increases TSLP and drives stromal TSLP-dependent ILC2 activation. **(A)** Schematic of the protocol for measuring tissue TSLP after feeding. Mice were fasted for 16 hr overnight (O/N) before oral gavage with 500 μl food slurry (refed) or water as volumetric control (sham). Tissues were harvested as indicated. **(B)** Percentage of TSLP-tdTomato^+^ cells among CD45^-^ cells in proximal jejunal small intestine lamina propria (siLP) by flow cytometric analysis after oral gavage at 2 hr. **(C)** ELISA of TSLP protein recovered from proximal jejunal tissue explants after oral gavage at 2 hr in *Pou2f3^-/-^* or WT mice. **(D)** Schematic of the protocol for measuring siLP ILC2 activation after feeding. Mice were fasted 16 hr overnight and given access to standard chow and water *ad libitum* (refed) or maintained on water control (fasted). Tissues were harvested 4 hr later. (**E**)(**F**) Percentage of IL-13 (Smart13)^+^ ILC2 among ILC2s in proximal jejunal LP in *Tslpr^+/+^*Red5Smart13 or *Tslpr^-/-^*Red5Smart13 mice after 16 hr fasting followed by 4 hr refeeding *ad libitum*. (**E**) Representative flow plots. (**F**) Quantitation across experiments. ILC2s were gated as Lin^-^CD45^+^IL-5 (Red5)^+^ cells. (**G**)(**H**) Percentage of IL-13 (Smart13)^+^ ILC2 among ILC2s in proximal jejunal LP 4 hr after refeeding *ad libitum* in overnight fasted *Tslp^fl/fl^Pdgfrɑ^CreERT2+^*Smart13 or littermate control *Pdgfrɑ^+/+^*Smart13 mice (ctrl) post-tamoxifen. (**G**) Representative flow plots. (**H**) Quantitation across experiments. ILC2s were gated on Lin^-^CD45^+^GATA3^+^ cells. **(I)** Percentage of IL-13 (Smart13)^+^ ILC2 among ILC2s in proximal jejunal LP in *Tslpr^fl/fl^Il5^Cre+^*(Red5)Smart13 mice after 16 hr fasting followed by 4 hr refeeding *ad libitum* or maintained on water control. ILC2s were gated as Lin^-^CD45^+^IL-5 (Red5)^+^ cells. All data are biological replicates, n≥3. Data are representative of at least two independent experiments. Error bars indicate samples mean± SEM.*P<0.03, **P<0.002, ***P<0.0002.

### GLP-2 mediates telocyte TSLP release and activation of resident ILC2s

Primarily absorbed by the prevalent enterocytes, constituents in food are sensed by enteroendocrine cells (EECs) that respond to luminal content by releasing peptide hormones that regulate nutrient uptake, metabolism, intestinal motility and satiety. Classically named by dominant hormones they release, EECs develop along vertically zonated villus trajectories through nodes of differentiation accompanied by activation of overlapping components of their peptide hormone repertoires (*21,22*). We screened published databases for intestinal telocyte and trophocyte expression of gut peptide hormone receptors that would enable crosstalk between nutrient-sensing EECs and these subepithelial structural cells. In both mouse and human datasets (*15,16,23,24*), intestinal telocytes expressed high levels of mRNA for the receptor for glucagon-like peptide-2 (GLP-2) with less consistent evidence for other gut peptide hormone receptors (Fig. 3A-C and fig. S3D-E).

**Fig. 3.**
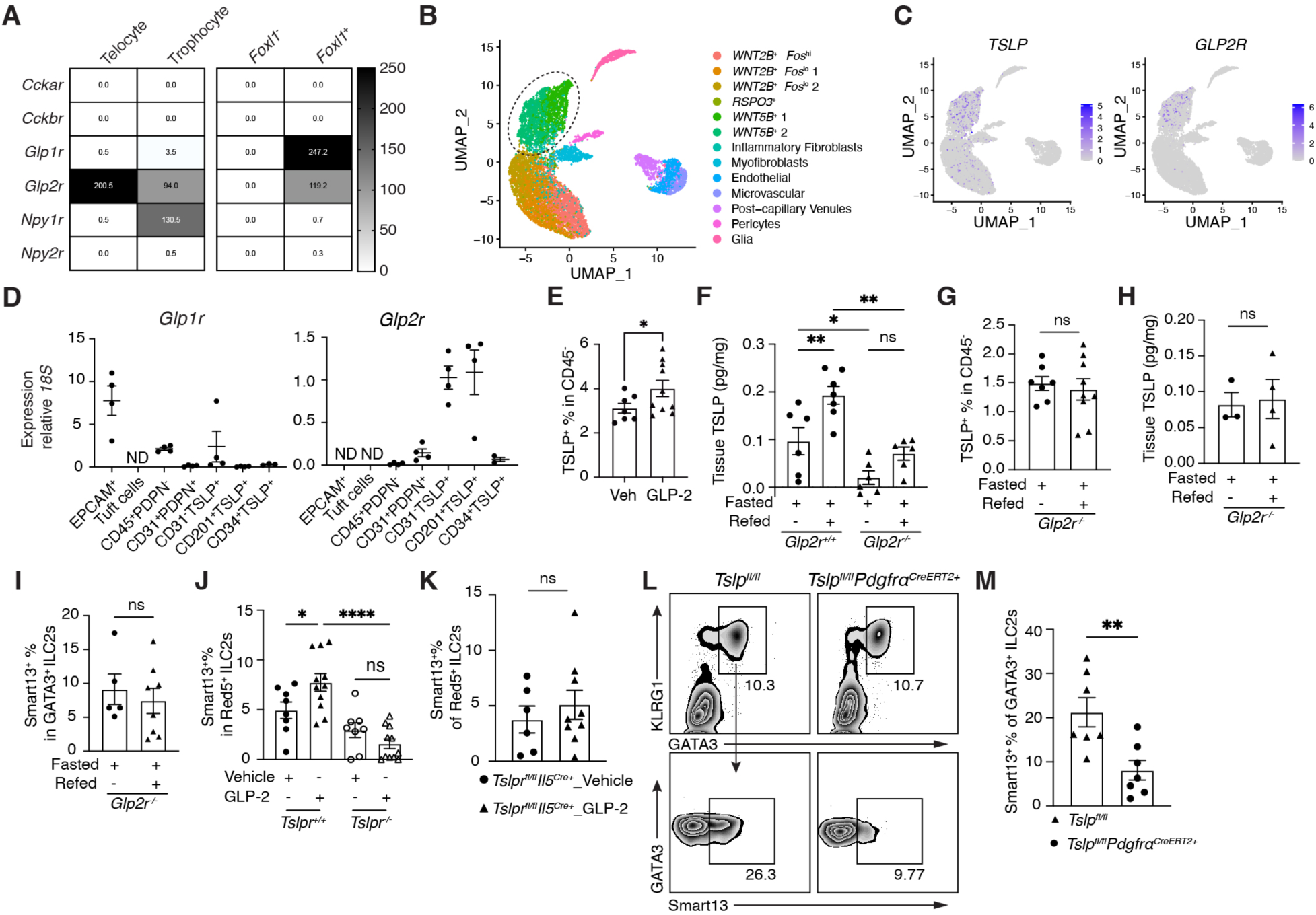
L cell GLP-2 drives GLP-2R- and TSLP-dependent ILC activation. (**A**) Heatmaps representing expression levels of gut hormone receptors in telocytes (*15*), trophocytes (*15*), *Foxl1*^+^ telocytes (*16*) and *Foxl1*^-^ stromal cells (*16*) from available mouse datasets. (**B**)(**C**) Single-cell analysis of an available human intestinal dataset (*23*). (**B**) UMAP representing cell clustering. (**C**) *TSLP* (**left**) and *GLP2R* (**right**) expression levels. (**D**) RT-qPCR analysis of *Glp1r* (**left**) and *Glp2r* (**right**) expression on purified gut EPCAM^+^ epithelial or TSLP-tdTomato^+^ stromal cells. (**E**) Percentage of TSLP-tdTomato^+^CD45^-^ cells in proximal jejunal lamina propria (LP) 2 hr after GLP-2[Gly2] injections as assessed by flow cytometric analysis. Veh, vehicle (**F**) ELISA for TSLP protein recovered from proximal jejunal tissue explants in *Glp2r*^-/-^ or wildtype (WT) mice 2 hr after GLP-2[Gly2] or vehicle control injections. (**G**) Flow cytometric analysis of percentage of total TSLP-tdTomato^+^CD45^-^ cells in proximal jejunal LP from fasted *Glp2r^-/-^*Flare-TSLP mice after oral food gavage at 2 hr. (**H**) ELISA for recovered TSLP protein in proximal jejunal homogenates from fasted *Glp2r^-/-^* mice after oral gavage at 2 hr. (**I**) Quantification of percentage of IL-13 (Smart13)^+^ ILC2 among proximal jejunal LP ILC2s from fasted *Glp2r^-/-^*Smart13 mice after 16 hr fasting followed by 4 hr refeeding *ad libitum* or maintained on water control. ILC2s were gated on Lin^-^CD45^+^GATA3^+^ cells. (**J**) Quantification of percentage of IL-13 (Smart13)^+^ ILC2s among proximal jejunal LP ILC2s in *Tslpr^+/+^*Red5Smart13 or *Tslpr^-/-^*Red5Smart13 mice after 3 daily subcutaneous (s.c.) injections of GLP-2[Gly2]. ILC2s were gated on Lin^-^CD45^+^IL-5(Red5)^+^ cells. (**K**) Quantification of percentage of IL-13 (Smart13)^+^ ILC2s among proximal jejunal LP ILC2s in *Tslpr^fl/fl^Il5^Cre+^* Smart13 after 3 daily subcutaneous (s.c.) injections of GLP-2[Gly2]. ILC2s were gated on Lin^-^CD45^+^IL-5(Red5)^+^ cells. (**L**) (**M**) Flow cytometric analysis of percentage of IL-13 (Smart13)^+^ ILC2 among proximal jejunal LP ILC2s in *Tslp^fl/fl^Pdgfrɑ^CreERT2+^*Smart13 or littermate *Pdgfrɑ^+/+^*Smart13 controls (ctrl) post-tamoxifen followed by 3 daily s.c. injections of GLP-2[Gly2]. (**L**) Representative flow plots. (**M**) Quantitation. ILC2s were gated as Lin^-^CD45^+^GATA3^+^ cells. All data are biological replicates, n≥3. Error bars indicate samples mean± SEM. *P<0.03, ****P<0.00001.

Although prominent in pancreatic glucagon-producing alpha cells, proglucagon is also highly expressed in subsets of small intestinal EECs, including L cells, which express tissue-specific convertases that post-translationally process proglucagon to the peptides GLP-1, GLP-2 and oxyntomodulin rather than glucagon (*25*, *26, 27*). In response to constituents in food, proglucagon-derived peptides are secreted into the basilar space, where activities of GLP-1 and GLP-2 are kept localized by high concentrations of the inactivating dipeptidyl peptidase, DPP-4. GLP-2 is intestinotrophic and promotes microvillus lengthening, increased vascular flow and an enhanced epithelial barrier, in part by stimulating release of insulin-like growth factor-1 (IGF-1) and epidermal growth factor (EGF) from subepithelial fibroblasts (*25*). We found *Glp1r* was expressed on both gut EPCAM^+^ epithelial and TSLP^+^ stromal cells whereas *Glp2r* was expressed on TSLP^+^ stromal cells and not detectable on epithelial cells (Fig. 3D), supporting an indirect role for GLP-2 on the gut epithelium. By crossing *Gcg^icre^*mice with *R26^lsl-YFP^* reporter mice, we could visualize preproglucagon^+^ cells by YFP expression. Using *Gcg*-YFP and Flare-TSLP dual reporter mice, we could observe L cells in proximity with TSLP^+^ telocytes after feeding (fig. S3F). Administration of stabilized GLP-2[Gly2] significantly increased the prevalence of TSLP-tdTomato^+^ cells in siLP as well as recovered TSLP protein in tissue explants as compared to animals treated with vehicle control or in mice lacking *Glp2r* (Fig. 3E-F, fig. S4A and S4E). The proportions of stromal or endothelial cells were not altered by GLP-2[Gly2] administration (fig. S4B), but there appeared to be more CD201^+^ telocytes within the stromal populations (fig. S4C-D). *In vitro* incubation of purified TSLP^+^ stromal cells with GLP-2[Gly2] resulted in increased antibody labeling of TSLP protein in the cells compared to vehicle control-treated cells (fig. S4F). In addition, feeding failed to increase the prevalence of TSLP-tdTomato^+^ cells in proximal siLP or amounts of TSLP protein recovered from intestinal tissue homogenates in fasted mice genetically lacking GLP-2R (Fig. 3G-H and fig. S4G). Activation of ILC2s was attenuated in *Glp2r*^-/-^ mice as assessed by IL-13 reporter (Smart13) expression (Fig. 3I). These results suggested that both TSLP production and ILC2 activation are downstream of GLP-2 in response to feeding. Further, injections of stabilized GLP-2[Gly2] led to increased frequencies of activated ILC2s that were attenuated in *Tslpr*^-/-^Red5Smart13 mice and *Tslpr^fl/fl^Il5^Cre^*Smart13 (Fig. 3J-K and fig. S4H-I), or *Tslp^fl/fl^Pdgfrɑ^CreERT2+^*Smart13 mice after tamoxifen treatment (Fig. 3L-M). Taken together, these data support a circuit linking activation of preproglucagon^+^ EECs with GLP-2-mediated stimulation of subepithelial telocytes to produce TSLP that activates resident ILC2s to produce IL-13.

### GLP-2 mediates ILC2-dependent amplification of the small intestinal tuft cell circuit

Activation of intestinal ILC2s to release IL-13 results in biased production of tuft cells from the transit-amplifying zone that is dependent on IL-4Rɑ expression on intestinal epithelial cells (*6,7*). In line with these data, administration of stabilized GLP-2[Gly2] resulted in increased numbers of small intestinal tuft cells that was significantly attenuated in *Tslpr*-, *Tslpr^fl/fl^Il5^Cre+^*, *Il13ra1^-^*^/-^ and *Il4ra^-^*^/-^ mice (Fig. 4A-B, fig. S5A-B). Further, the increase in tuft cells was attenuated in *Il4ra^fl/fl^*Vil1^Cre^ mice, supporting a role for epithelial IL-4Rɑ in activating this circuit (Fig. 4B and fig. S5A). To further confirm by an independent approach that EECs can activate the ILC2-tuft cell circuit through an activated G protein-coupled receptor (GPCR) pathway, we used triple mutant *Vil1^Flp^Cck^Cre^R26^Dual-hM3Dq^* mice with epithelial lineage-restricted expression of activating Designer Receptors Exclusively Activated by Designer Drugs (DREADDs) on CCK-lineage-restricted intestinal epithelia, which includes preproglucagon^+^ EECs that secrete GLP-2 (*22*) (Fig. 4C, fig. S5C). After administration of agonist clozapine N-oxide (CNO), tuft cells were increased in the small intestine, confirming amplification of the epithelial tuft cell circuit downstream of EEC activation (Fig. 4D-F and fig. S5D-E).

**Fig. 4.**
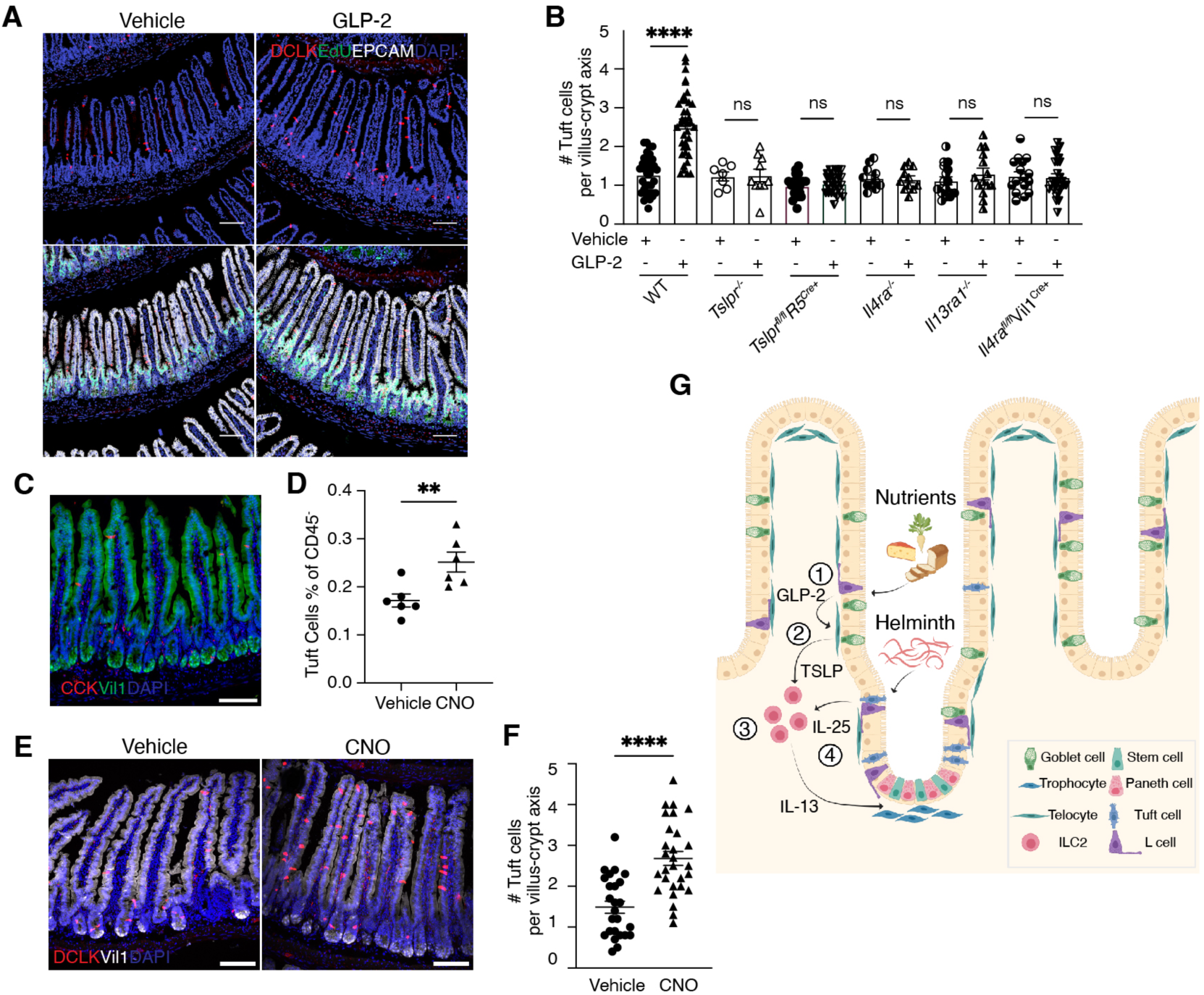
GLP-2 drives ILC2-dependent amplification of the tuft cell cycle. (**A**) Representative imaging of tuft cells in jejunum after 3 daily injections of GLP-2[Gly2]. 5-ethynyl-2’-deoxyuridine (EdU) was injected the night before tissue harvest. Green, EdU; Red, DCLK; White, EPCAM; Blue, DAPI. 20x objective. Scale bar, 100 µm. (**B**) Quantification of jejunal tuft cells after 3 daily GLP-2[Gly2] injections in WT, *Tslpr*^-/-^, *Tslpr^fl/fl^Il5^Cre+^*, *Il4ra*^-/-^, *Il13ra*^-/-^ and *Il4ra^fl/fl^*Vil1^Cre^ mice. For tuft cell quantification, images were acquired using 20x objective and total DCLK^+^EPCAM^+^ cells in each field were counted and normalized by total number of villus-crypt axes in the field. Each dot represents an imaging field; data pooled from 3-5 mice. (**C**) Representative imaging showing CCK^+^ EEC cells in *Vil1^Flp^Cck^Cre^R26^Dual-hM3Dq^* mice. Red, *Cck*-mCherry; Green, Vil1-GFP; Blue, DAPI. 20x objective. Scale bar, 100 µm. (**D**) Percentage of tuft cells among the CD45^-^ epithelial fraction after administration of CNO or vehicle control. (**E**) (**F**) Quantification of jejunal tuft cells after administration of CNO or vehicle control. (**E**) Representative imaging of tuft cells. Red, DCLK; White, *Vil1*-GFP; Blue, DAPI. 20x objective. Scale bar, 100 µm. (**F**) Average tuft cell numbers in each villus-crypt unit. Each dot represents an imaging field; data pooled from 4 mice. (**G**) Proposed model. GLP-2 from L cells (**1**) stimulates telocytes to release TSLP (**2**) that activates ILC2s (**3**), thus linking nutrient ingestion and IL-13-mediated amplification of epithelial tuft cells (**4**). *Nippostrongylus brasiliensis* short circuit the physiologic pathway. Created using BioRender. All data are biological replicates, n≥3. Error bars indicate samples mean± SEM. *P<0.03, **P<0.002, ***P<0.0002.

## Discussion

Constituents of many tissues often in relation to stem cell niches (*28*), telocytes and trophocytes are structural subepithelial fibroblasts extending from the crypts to the villus tips whose arborizing dendrites establish the critical WNT and BMP counter-gradients that maintain stem cells and control epithelial differentiation during villus transit (*15,16,29*). As assessed in Flare-TSLP reporter mice, villus telocytes and crypt-associated trophocytes are the major cells that constitutively express TSLP, an alarmin linked with activation of type 2 immune cells, including ILC2s, and support findings identifying long-lived structural cells as hubs for tissue immunity (*30*). Across small intestine, TSLP expression was prominent in ‘tip’ telocytes, or VTTs, specialized *Lgr5*^+^ fibroblasts that control epithelial differentiation at villus apices, a localization positioning these cells to relay information from epithelia regarding intestinal constituents (*17*).

Although activation of ILC2s by food intake was noted (*19*), the pathway by which information regarding luminal nutritional content is relayed through epithelia to resident ILC2s was unknown. EECs sense nutrient content and secrete peptide hormones and neurotransmitters that collectively orchestrate intestinal motility, absorption, and integration of neuronal circuitry controlling feeding, satiety and positive consumptive or negative aversive reactions (*31*). Sophisticated fate-mapping and lineage-marking studies revealed EECs have a longer half-life than enterocytes during villus ascent and are vertically zonated such that peptide hormones are sequentially expressed in over-lapping patterns among EEC subsets to optimize small intestinal function (*21,22,32*). By scanning available databases and our own samples, we verified the high expression of receptors on subepithelial telocytes for GLP-2, a proglucagon-derived peptide previously shown to promote release of epithelial growth factors from subepithelial fibroblasts (*33*). Although GLP-2 deletion does not impact small intestine differentiation, it is required for recovery from intestinal atrophy after fasting or injury (*34–36*), underpinning the success of stabilized GLP-2 receptor agonists used therapeutically in humans to increase small intestine growth or prevent atrophy following parenteral feeding (*26*). Although effects of GLP-2 receptor agonists are reversible after cessation of therapy (*37*), the striking intestinotrophic effects of model helminth and protist infections prompted us to query whether this physiologic pathway shared mechanisms with the intestinal response to these organisms and reflecting the integration of immunity with tissue homeostasis (*38*).

As we show, GLP-2 stimulates telocytes to release TSLP, an alarmin that activates ILC2s, thus linking nutrient ingestion and IL-13-mediated amplification of epithelial tuft cells. Tuft cells are specialized chemosensory cells that express a variety of GPCRs that share signal transduction recognition with type 2 taste receptors and provide feedback amplification of the ILC2 - tuft cell circuit by release of a second alarmin, IL-25 (*39*). As recently suggested, such a system may be part of a regulated food quality control circuit by increasing detection of constituents in ingested cargo that attach positive or negative associations on content that has passed more proximal surveillance mechanisms (*40*). In evolution, eating may have been associated with dispersed periods involving ingestion of large kills over several days that would favor inducible mechanisms involving enhanced sensitivity for potentially toxic products of decay. Although nutrients stimulate EECs and indirectly activate the tuft cell circuit via GLP-2, intestinal protists and helminths directly stimulate tuft cell GPCRs and sustain the circuit (*9,41,42*). By hijacking a physiologic circuit linked with feeding, endemic parasites establish a reproductive niche protected from super-infection by other pathogens while sustaining the host metabolic state (*9,43,44*), a condition supported by increased attentiveness to ingested constituents (Fig. 4G).

## Acknowledgments

The authors thank A. Ma and A. B. Molofsky for constructive review of the manuscript, M. S. Anderson, Z. A. Knight, S. F. Ziegler, F. de Sauvage, and O. D. Klein for mouse lines, current and former members of the Locksley laboratory for comments, UCSF core facility for technical support, M. Conseco for mouse colony support and A. Luong for administrative support.

## Funding

National Institutes of Health grant AI026918 and HL107202 (RML) Howard Hughes Medical Institute (RML)

Sandler Basic Asthma Research Center at University of California, San Francisco (RML)

Canadian Institute Health Research (CIHR) grant 154321(DJD) Cancer Research Institute Fellowship (CL)

## Author contributions

CL conceived the study, designed and performed experiments, analyzed and interpreted the data, and wrote the manuscript. HEL generated the Flare-TSLP reporter mice and participated in experiments. MJ and ZW participated in experiments. DJD provided key material. RML conceived and directed the studies with CL, analyzed and interpreted data, and wrote the manuscript with CL with input from the co-authors.

## Data and materials availability

All data are available in the main text or the supplementary materials.

## Supplementary Materials for

### Materials and Methods

#### Mice

Mice were C57BL/6J purchased from the Jackson laboratory (JAX #000664) or mutant alleles backcrossed >8 generations to C57BL/6J. Red5 or B6(C)-*Il5^tm1.1(icre)Lky^*/J (*19*), Smart13 or B6.129S4(C)-*Il13^tm2.1Lky^*/J (*10*), *Tslpr^-/-^* (*45*), backcrossed *Il4ra*^-/-^ or BALB/c-*Il4ra^tm1Sz^*/J (#003514) (*6*), and *Il13ra1*^-/-^ (*46*) mice generated or obtained by this lab have been described. Vil1^Cre^ or B6.Cg-Tg(Vil1-cre)997Gum/J (JAX #004586), *Pdgfra^CreERT2^* or B6.129S-*Pdgfra^tm1.1(cre/ERT2)Blh^*/J (JAX #032770), *Gcg^iCre^* or B6;129S-*Gcg^tm1.1(icre)Gkg^*/J (JAX #030663) and *R26^lsl-EYFP^* or B6.129X1-*Gt(ROSA)26Sor^tm1(EYFP)Cos^*/J (JAX #006148) were purchased from the Jackson laboratory. C57BL/6 *Pou2f3*^-/-^ (re-derived from C57BL/6NTac *Pou2f3^tm1.1(KOMP)Vlcg^*) mice (*9*) were provided by M. S. Anderson. C57BL/6 *Tslpr^fl/fl^* mice (*47*) were provided by S. F. Ziegler. C57BL/6 *Glp2r^-/-^* mice (*48*) were provided by D. Drucker. *Lgr5*-eGFP mice (*49*) were provided from Genentech by F. de Sauvage and O. D. Klein. Triple-mutant *Vil1^Flp^ Cck^Cre^* (JAX #012706) *R26^Dual-hM3Dq^*(JAX #026942) mice (*22*) were provided by Z. A. Knight.

Flare-TSLP mice were generated by homologous gene targeting in C57BL/6 embryonic stem cells based on a reporter cassette used to target the *Il25* locus (*6*). Briefly, a 2.34 kb 3’ homologous arm in the 3’UTR of *Tslp* was amplified from C57BL/6 genomic DNA and a 3.79 kb fragment from exon 1 to the end of coding sequence of exon 5 was amplified to serve as the 5’ homologous arm. A 5’*loxP* site was inserted 187 bp upstream of exon 4 within the modified 5’ arm. The reporter cassette encoded (5’ to 3’) a *loxP* site (3’*loxP* in final construct), encephalomyocarditis virus IRES element, tandem RFP or tdTomato, bovine growth hormone poly(A) signal, and an frt-flanked neomycin selection cassette, and was subcloned between the modified 5’ and 3’ homologous arms with correct orientation using the multiple cloning site of a basal targeting vector, pKO915-DT (Lexicon Genetics). After linearization by NotI the reporter cassette was electroporated into C57BL/6 ES cells. Following growth on irradiated mouse embryonic fibroblast feeders and selection in G418-containing media, neomycin-resistant clones were screened for correct 5’ and 3’ homologous recombination by long-range PCR and further confirmed for retention of the 5’*loxP* site. One clone was injected into albino C57BL/6 blastocysts to generate chimeras and males coat with the highest black-to-white coat color ratio were bred with B6.129S4-*Gt(ROSA)26Sor^tm1(FLP1)Dym^*/RainJ females (JAX#009086) to excise the neomycin resistance cassette. Offspring with germline transmission and confirmed neo^r^-deleted *Flare-Tslp* allele were backcrossed to C57BL/6 mice to cross out the FLP1 allele. *Flare-Tslp* genotyping was done using primers 5’*loxP fw*: 5’- GGAACGAAGTTGAAACACCACGACC-3’and 5’*loxP rev*: 5’-GGGGATGGGGATAGGAGGGAAAGAC-3’, yielding a 277 bp mutant or a 243 bp wildtype band. After Cre-*loxP*-mediated deletion, the floxed-out *Tslp* allele can be detected by PCR using primers 5’*loxP fw* (as above) and *IRES rev*: 5’-TTGTTGAATACGCTTGAGGAGAGCC-3’, yielding a 596 bp band while a non-recombined floxed band is 2878 bp.

Mice were maintained in the University of California, San Francisco (UCSF) specific pathogen-free animal facility in accordance with the guidelines established by the Institutional Animal Care and Use Committee and Laboratory Animal Resource Center. Unless otherwise specified, mice were kept on 12 hr light-dark cycles and *ad libitum* water and chow (PicoLab 5053). Pooled experiments included female and male mice. All experimental procedures were approved by the Laboratory Animal Resource Center at the UCSF.

#### Antibodies

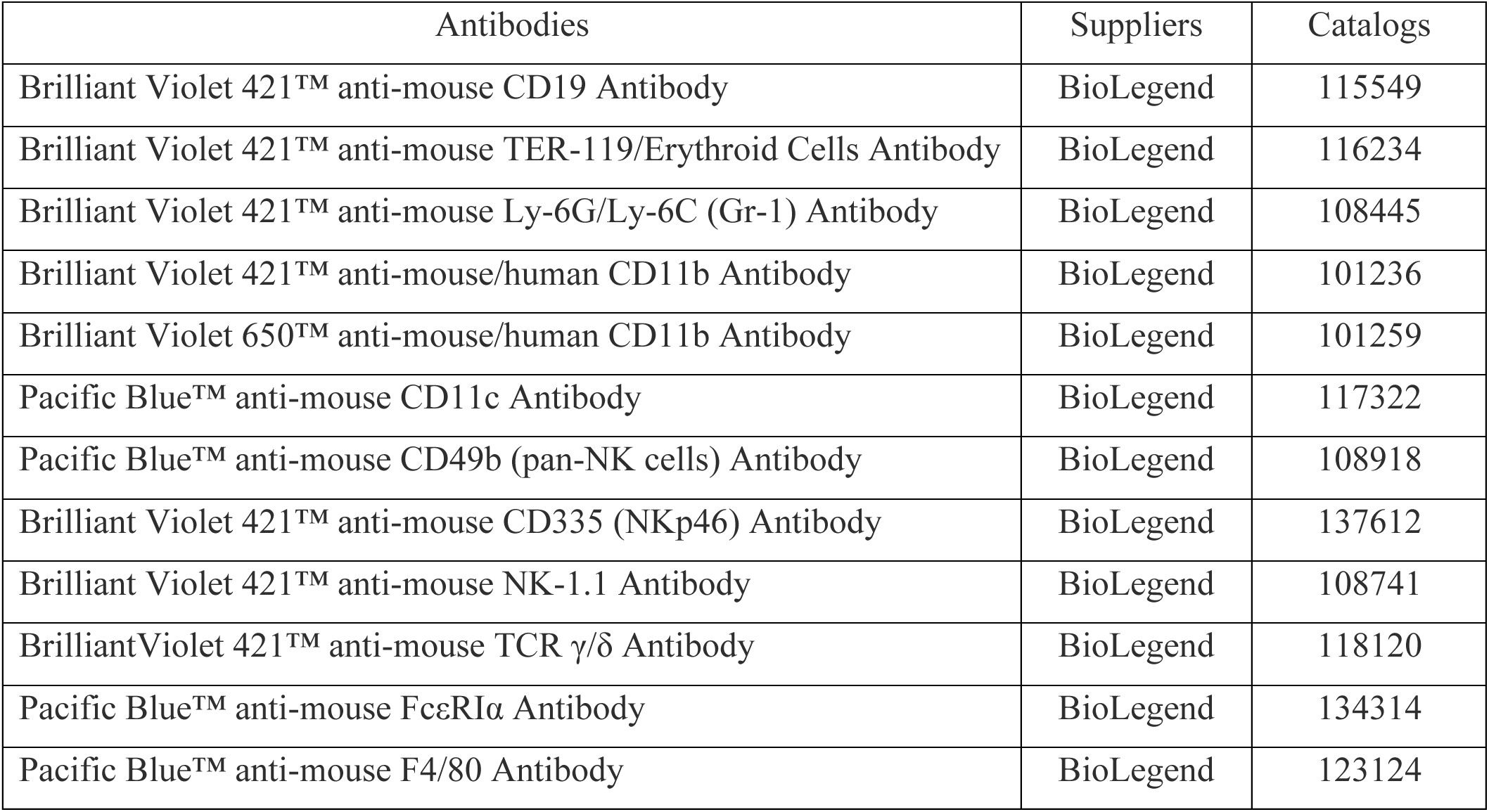
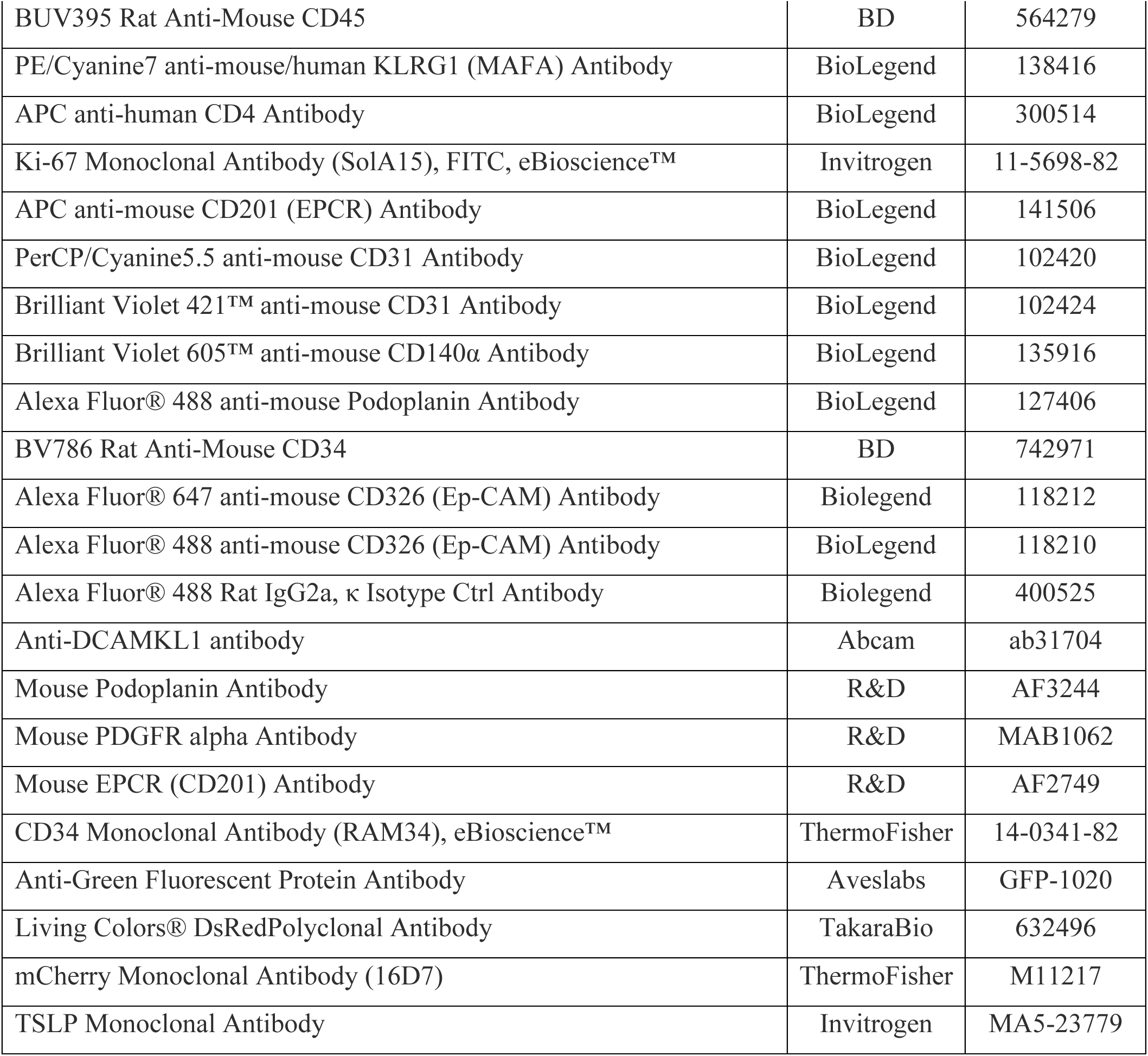

#### TSLP antibody labeling

Mouse TSLP antibody for flow cytometric analysis was labeled using Alexa Fluor™ 488 Antibody Labeling Kit (ThermoFisher, Cat# A20181) following the manufacture protocol.

#### Flow cytometry and fluorescence-activated cell sorting (FACS)

Small intestine tissues were harvested at indicated times and single-cell suspensions of epithelium or lamina propria were obtained. In brief, 5-6 cm proximal jejunum or distal ileum were harvested and flushed with ice-cold PBS. After removing the Peyer’s patches, gut tissues were opened and washed with PBS to remove lumen content and mucus. Segments were incubated while rocking for 15 min at 37 °C in 20 ml HBSS (Ca^2+^- and Mg^2+^-free) containing 5% FCS, 10 mM HEPES (Sigma, Cat#H3537), 10 mM EDTA (Corning, Cat#46-034-CI) and 10 mM dithiothreitol (DTT; Sigma, Cat#D0632). Tissue segments were vortexed vigorously for 30 s and supernatants collected and centrifuged. The pelleted epithelial fraction was washed once with PBS before staining for flow cytometry. The rest of the tissue segments were incubated a second time while rocking for 15 min at 37 °C in 10 ml of the same HBSS/FBS/EDTA/DTT media and then vortexed for 30 sec. The tissue segments were transferred to a new tube and incubated while rocking for 5 min at 37 °C in 20 ml HBSS (with Ca^2+^ and Mg^2+^) containing 2% FCS and 10 mM HEPES before transferring to new tubes containing 5 ml HBSS (with Ca^2+^ and Mg^2+^), 3% FCS, 10 mM HEPES, 30 µg/ml DNase (Roche, Cat#10104159001) and 0.1 unit/ml Liberase TM (Roche, Cat#5401127001) in C tubes (Miltenyi Biotec, Cat# 130-096-334).

Samples were processed with gentleMax Octo Dissociator program “intestine”. Lamina propria cells were pelleted, washed and filtered with 70 µM cell strainers before staining for flow cytometry. Standard surface and/or intracellular staining protocols were used followed by DAPI staining for dead cell exclusion. Samples were processed using a BD LSRFortessa™ (Bectin Dickinson) flow cytometer and analyzed using FlowJo software. Cell sorting experiments were performed on MoFlo (Beckman Coulter) or BD FACSAria™ (Bectin Dickinson).

#### Tissue immunofluorescence

Tissues were fixed in 4% paraformaldehyde (PFA; Electron, Cat#15710) for 2 hr at room temperature (RT) or longer at 4°C, washed 3 times in PBS and incubated in 30% (w/v) sucrose-PBS solution overnight at 4°C. Tissues were embedded in Optimal Cutting Temperature Compound (OCT; VWR, Cat#25608-930) on dry ice and stored at −80 °C. Skin samples were cut into 1 cm x 2 cm pieces prior to fixation. Samples from proximal jejunum, mid gut, and distal ileum (∼10 cm) were cut and flushed with PBS before fixation and coiled into “Swiss rolls” before embedding. OCT embedded tissues were cut into 10-12 µm regular sections or 100 µm thick sections using Cryostat (Leica CM3050 S) and stored at −80 °C. Sections were incubated with 0.02% Triton X-100 (Sigma, Cat#X100) in TNB buffer (Tris-NaCl-blocking buffer: 0.1M Tris-HCl, PH 7.5; 0.15M NaCl; Blocking reagent, Akoya Biosciences, Cat#FP1020) for permeabilization followed by blocking with 5% serum matching the source of the secondary antibodies for 30 min at RT. After 2 hr incubation with primary antibodies at RT, sections were washed three times in PBS and incubated with secondary antibodies for 1 hr at RT. Sections were incubated with DAPI solution for 5 min and washed three times in PBS before mounting. 5-ethynyl-2’-deoxyuridine (EdU) staining was performed using Click-IT Plus EdU imaging kit (Invitrogen, Cat#C10637) following manufacture manual. Stained samples were visualized using a Nikon A1R confocal microscope with 20x objective and analyzed using NIS Element and ImageJ software. Thick section staining protocols were as described (*50*) and imaging was processed using Imaris software - 3D analysis workstation for Cell Biology Research (Oxford Instruments).

#### TSLP ELISA from intestinal homogenates and explants

Tissue samples from proximal jejunum (2 cm) or distal ileum (3 cm) were weighed and harvested in RIPA (ThermoFisher, Cat. 89900) buffer containing proteinase inhibitors in M tubes (Miltenyi Biotec, Cat. 130-096-335) followed by dissociation with gentleMax Octo Dissociator program “protein 1”. Supernatants were collected following brief centrifugation and stored at −80 °C. TSLP ELISA was performed using the Biolegend Deluxe TSLP ELISA kit (BioLegend, Cat. 434104) with undiluted homogenates and protein concentrations were normalized by tissue weight of the starting samples.

#### Drug Treatments

MC903 (Calcipotriol, Tocris Biosciences, Cat. 2700, 100 µM working concentration in ethanol) was applied topically on shaved back skin using 60 µl per mouse daily for 7 consecutive days (*14*). Stabilized human GLP-2[Gly2] (teduglutide, Echelon Bioscience, Cat. 471-21) was given at 10 µg/mouse subcutaneously in 100 µl DMSO-PBS solution consecutively for 3 days or as indicated at 25 µg/mouse i.p. 2 hrs prior to harvesting tissues. Tamoxifen (Sigma, Cat. T5648) was administrated at 3 mg/mouse or 100 mg/Kg in 100 µl corn oil i.p. daily for 7 days; mice were rested for 3-5 days before experimental use. Alternatively, tamoxifen diet (Inotive, Cat. TD.130858) was given to mice *ad libitum* for at least 4 wks prior to experimental use. Clozapine N-oxide (CNO; Cayman, Cat. 16882) was given 5 mg/kg i.p. for 3 consecutive days. EdU (Sigma, Cat. 900584) was given i.p. at 1 mg/mouse in 100 µl PBS 24 hrs prior to harvesting tissue.

#### Nippostrongylus brasiliensis

*N. brasiliensis* was maintained in rats and purified as described (*6*). Mice were injected s.c. with 500 purified L3 larvae and small intestines were harvested on day 5, 7, and 12 post infection (d.p.i) for analysis of epithelial tuft cells as described (*6*).

#### Fasting - feeding protocols

For measuring small intestinal TSLP, designated mice were fasted 16 hr overnight before oral gavage with 500 µl food slurry (modified powdered diet using formula TekladTD.88232 with comparable nutrient distribution as standard chow diet) in water containing 1.965 Kcal for ∼1% daily caloric intake) or water as a volumetric control. For small intestinal ILC2 activation, designated mice were fasted overnight for 16 hr before restoring access to standard chow diet (PicoLab 5058) and water *ad libitum* or maintaining on water. Tissues were harvested at indicated time points. Overnight fasting followed by water gavage or access *ad libitum* was designated the “Fasted” control group; overnight fasting followed by food gavage or access *ad libitum* was designated the “Fasted+refed” group.

#### Tuft cell quantifications

For tuft cell quantification, images were acquired using 20x objective and total DCLK^+^EPCAM^+^ cells in each field were counted and normalized by total number of villus-crypt axes in the field.

#### Quantitative polymerase chain reaction (qPCR) and gene expression analysis

For analysis of designated populations, cells were purified by FACS directly into lysis buffer followed by RNA extraction using the Qiagen kit (Cat# 74034). cDNA was synthesized using Invitrogen SuperScript IV VILO kit (Invitrogen, Cat# 11754250). qPCR reactions were performed using Taqman probes (Invitrogen) and Taqman gene expression master mix (Invitrogen, Cat# 4369016). Transcripts were normalized to *18S* expression. Probe information as follows:

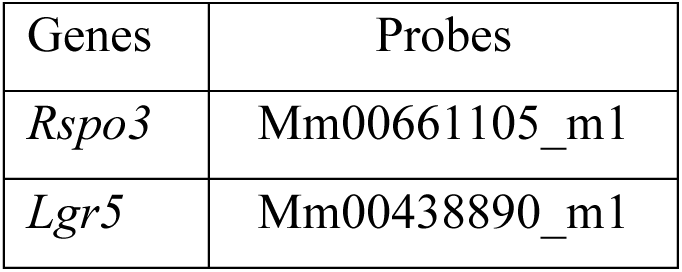
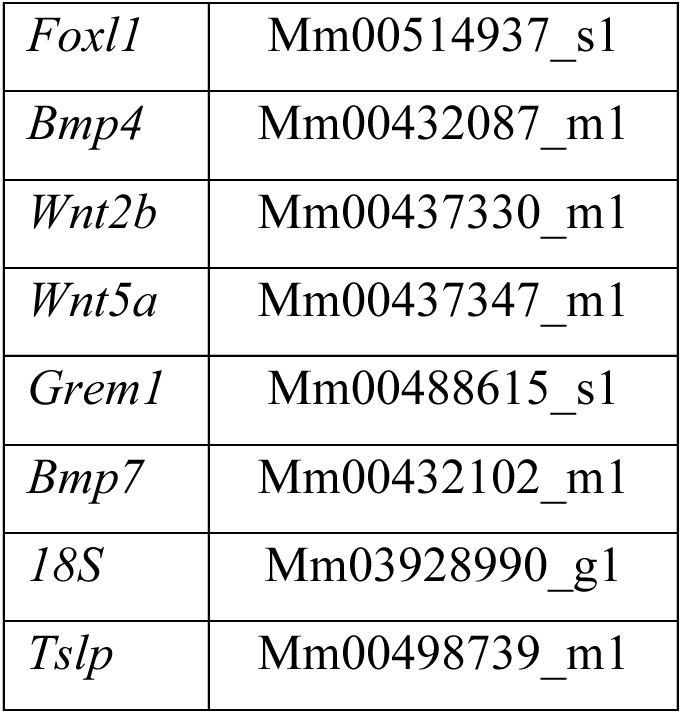

#### Single-cell RNA sequencing data analysis

Available human datasets (raw or pre-processed data) (*23*,*24*) were obtained from the single-cell portal of the Broad Institute (https://singlecell.broadinstitute.org). Data were re-analyzed and plotted using Seurat package in R program.

#### Statistics

Quantitative data were represented as mean ± standard error of the mean (SEM) of at least duplicate biological replicates. Student t test was used for analyses of difference between 2 groups. Analysis of variance (ANOVA) was used for analyzing experiments with >2 groups adjusted for multiple comparisons. Significance level was set at α = 0.05. Statistical analysis was performed using Prism 9 (GraphPad Software). *P<0.03, **P<0.002, ***P<0.0002, ****P<0.00001.

---

**Fig. S1.**
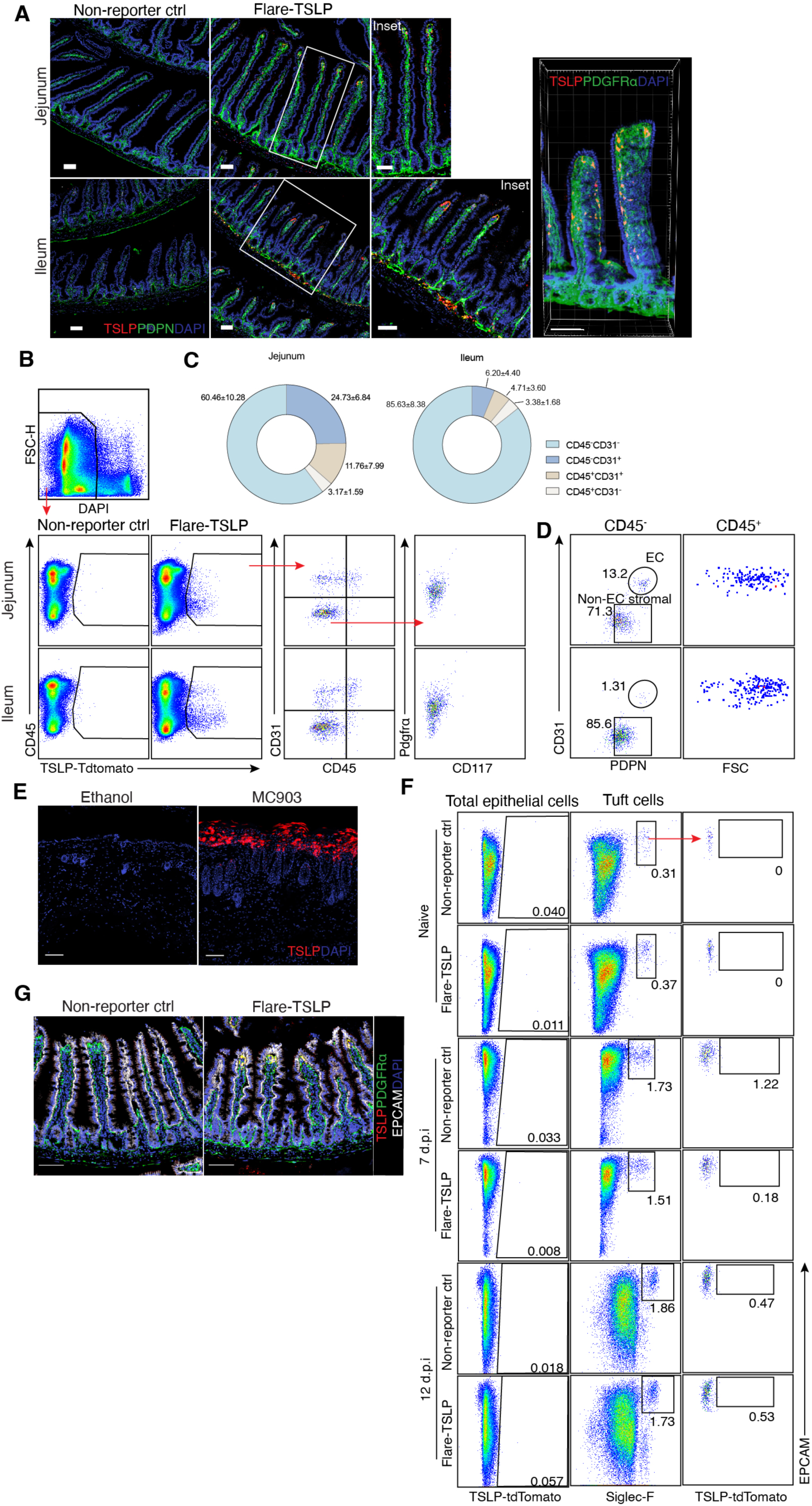
Characterization of Flare-TSLP reporter mice. (**A**) Representative imaging showing jejunum (**top**) and ileum (**bottom**) of Flare-TSLP mice or non-reporter control mice. Red, TSLP-tdTomato; Green, PDPN; Blue, DAPI. 20x objective. Scale bar, 50 µm. **Right panel**, max intensity projection of three - dimensional imaging showing jejunual tissues from Flare-TSLP mice ; scale bar, 100 µm. (**B**)-(**D**) Flow cytometric analysis of CD45^+^, endothelial and non-endothelial stromal populations in jejunal and ileal lamina propria in Flare-TSLP or non-reporter control (ctrl) mice. (**B**) CD45^+^ cells were CD31^+^ specialized endothelial cells. CD45^-^CD31^-^ stromal cells were PDGFRɑ^+^cKit^-^ (non-Cajal cells, which are cKit^+^). (**C**) Proportions of CD45^+^CD31^+^, CD45^+^CD31^-^, CD45^-^CD31^+^, and CD45^-^CD31^-^ cells in TSLP-tdTomato^+^ cells represented by pie charts. (**D**) TSLP-tdTomato^+^CD45^-^CD31^+^ cells were PDPN^+^ consistent with lymphatic endothelial cells. TSLP-tdTomato^+^CD45^+^ cells are CD31^+^ FSC^hi^ large cells consistent with a non-hematopoietic lineage. (**E**) Representative imaging showing back skin of Flare-TSLP mice treated with ethanol or MC903 for 7 days. Red, TSLP-tdTomato; Blue, DAPI. 20x objective. Scale bar, 100 µm. (**F**) Flow cytometric analysis of epithelial TSLP expression in proximal small intestine in naive Flare-TSLP or non-reporter mice or on day 7 and 12 post *N. brasiliensis* infection. d.p.i., day post infection. (**G**) Representative imaging of jejunum in Flare-TSLP or non-reporter mice on day 5 after *N. brasiliensis* infection. Red, TSLP-tdTomato; Green, PDGFRɑ; White, EPCAM; Blue, DAPI. 20x objective. Scale bar, 100 µm. All data are biological replicates, n≥3. Data are representative of at least two independent experiments.

**Fig. S2.**
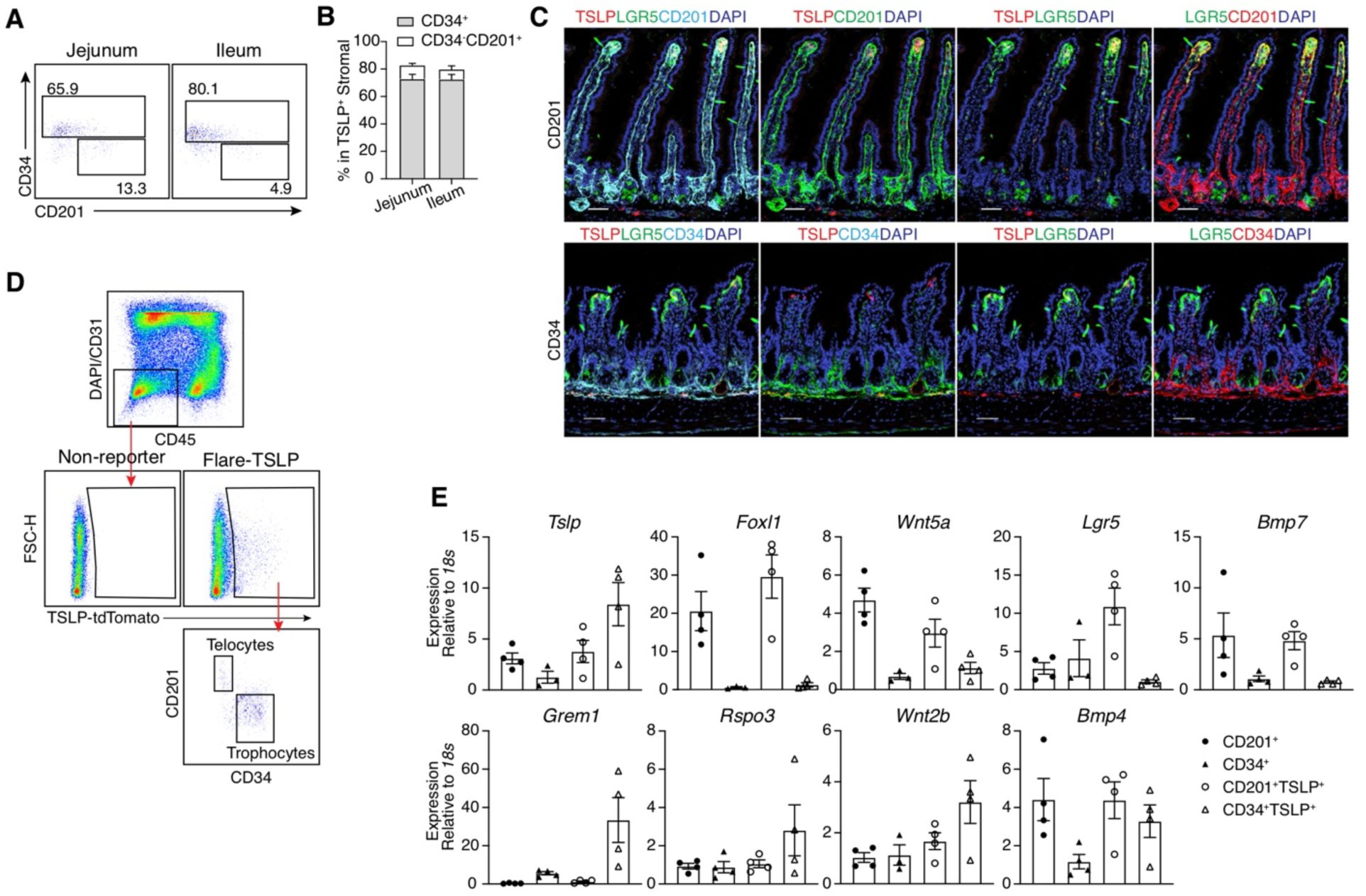
Analysis of CD201^+^ telocytes and CD34^+^ trophocytes. (**A**) (**B**) CD201 and CD34 staining of TSLP-tdTomato^+^ cells in jejunal and ileal lamina propria (LP) of Flare-TSLP mice. (**A**) Representative flow cytometric analysis. (**B**) Quantification of percentage of CD34^+^ and CD34^-^CD201^+^ in TSLP-tdTomato^+^ populations. (**C**) Representative imaging of CD201 and CD34 expression in proximal and distal small intestine of Flare-TSLP; *Lgr5*-eGFP dual reporter mice. Scale bar, 50 µm. (**D**) Gating strategy for sorting of TSLP-tdTomato^+^CD201^+^CD31^-^, and TSLP-tdTomato^+^ CD34^+^CD31^-^ cells from small intestinal LP (siLP, whole tissue) of Flare-TSLP mice. (**E**) RT-qPCR analysis of purified TSLP-tdTomato^+^CD201^+^CD31^-^ (telocytes), and TSLP-tdTomato^+^ CD34^+^CD31^-^ (trophocytes), CD201^+^CD31^-^, and CD34^+^CD31^-^ stromal cells from siLP (whole tissue) of Flare-TSLP mice. All data are biological replicates, n≥3. Data are representative of at least two independent experiments. Error bars indicate samples mean± SEM.

**Fig. S3.**
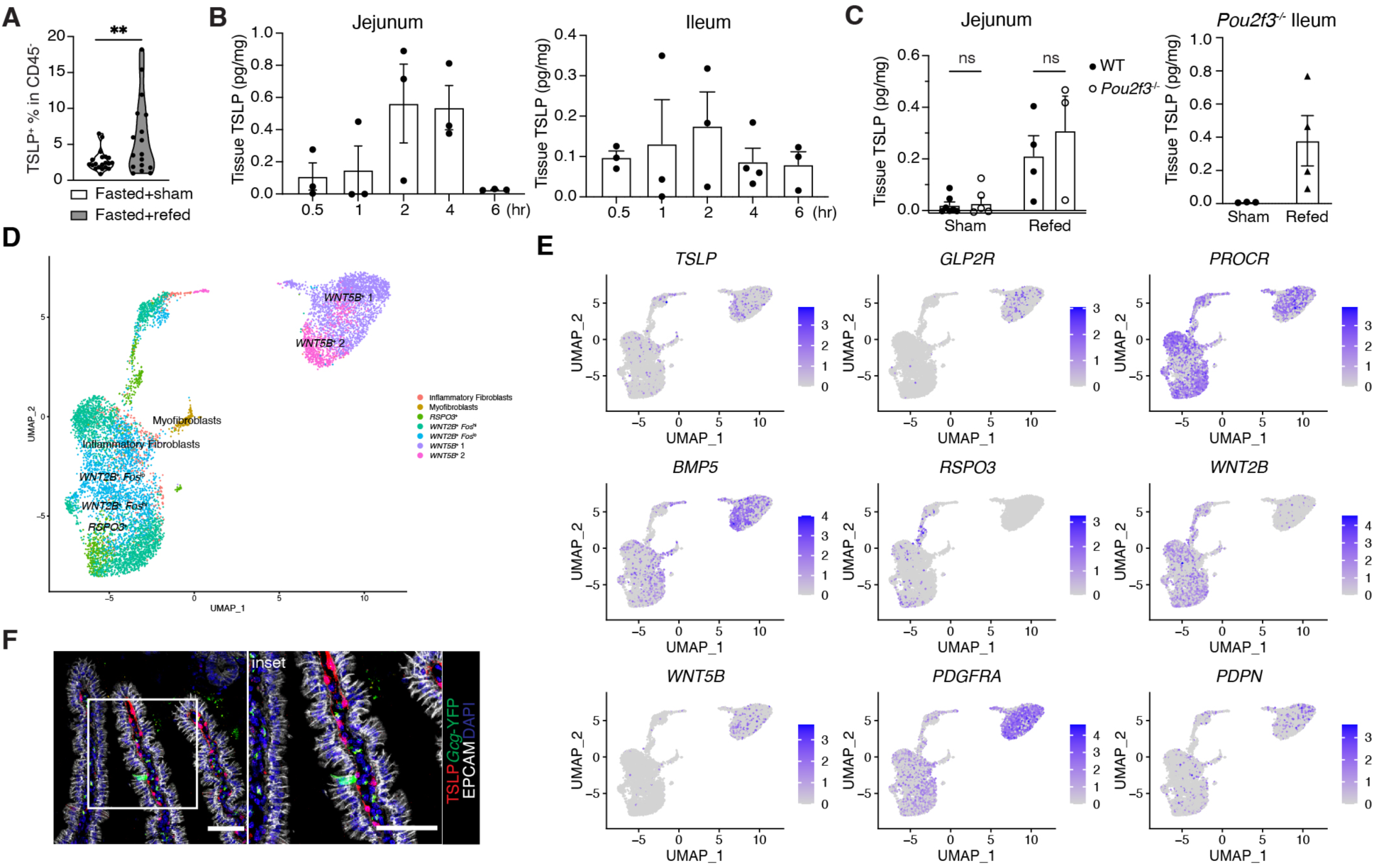
Feeding increases TSLP and drives stromal TSLP-dependent ILC2 activation. (**A**) Percentage of TSLP-tdTomato^+^ cells among CD45^-^ cells in ileal lamina propria by flow cytometric analysis in fasted mice at 2 hr after oral gavage of food (refed) or water as volumetric control (sham). (**B**) ELISA of TSLP protein recovered from proximal jejunal (**left**) and distal ileal (**right**) tissue homogenates after oral gavage of food at designated times in fasted wild type (WT) mice. (B) ELISA of TSLP protein recovered from jejunum and distal ileal tissue homogenates from fasted *Pou2f3*^-/-^ mice at 2 hr after oral food gavage. **(D)(E)** Single-cell analysis of a human intestinal dataset (*24*). (**D**) UMAP representing cell clustering. (**E**) Gene expression levels of *TSLP*, *GLP2R*, *FOXL1*, *GREM1*, *CD34*, *PROCR*, *BMP5*, *RSPO3*, *WNT2B*, *WNT5B*, *PDGFRA*, and *PDPN*. (**F**) Representative imaging showing a preproglucagon^+^ EEC (L cell) and TSLP-tdTomato^+^ telocyte in proximity in the jejunum of *Gcg*-YFP; Flare-TSLP dual reporter mice after 16 hr overnight fasting followed by refeeding *ad libitum* for 4 hr. Red, TSLP-tdTomato; Green, *Gcg*-YFP; White, EPCAM; Blue, DAPI. 20x objective. Scale bar, 50 µm. All data are biological replicates, n≥3. Data are representative of at least two independent experiments. Error bars indicate samples mean± SEM. **P<0.002.

**Fig. S4.**
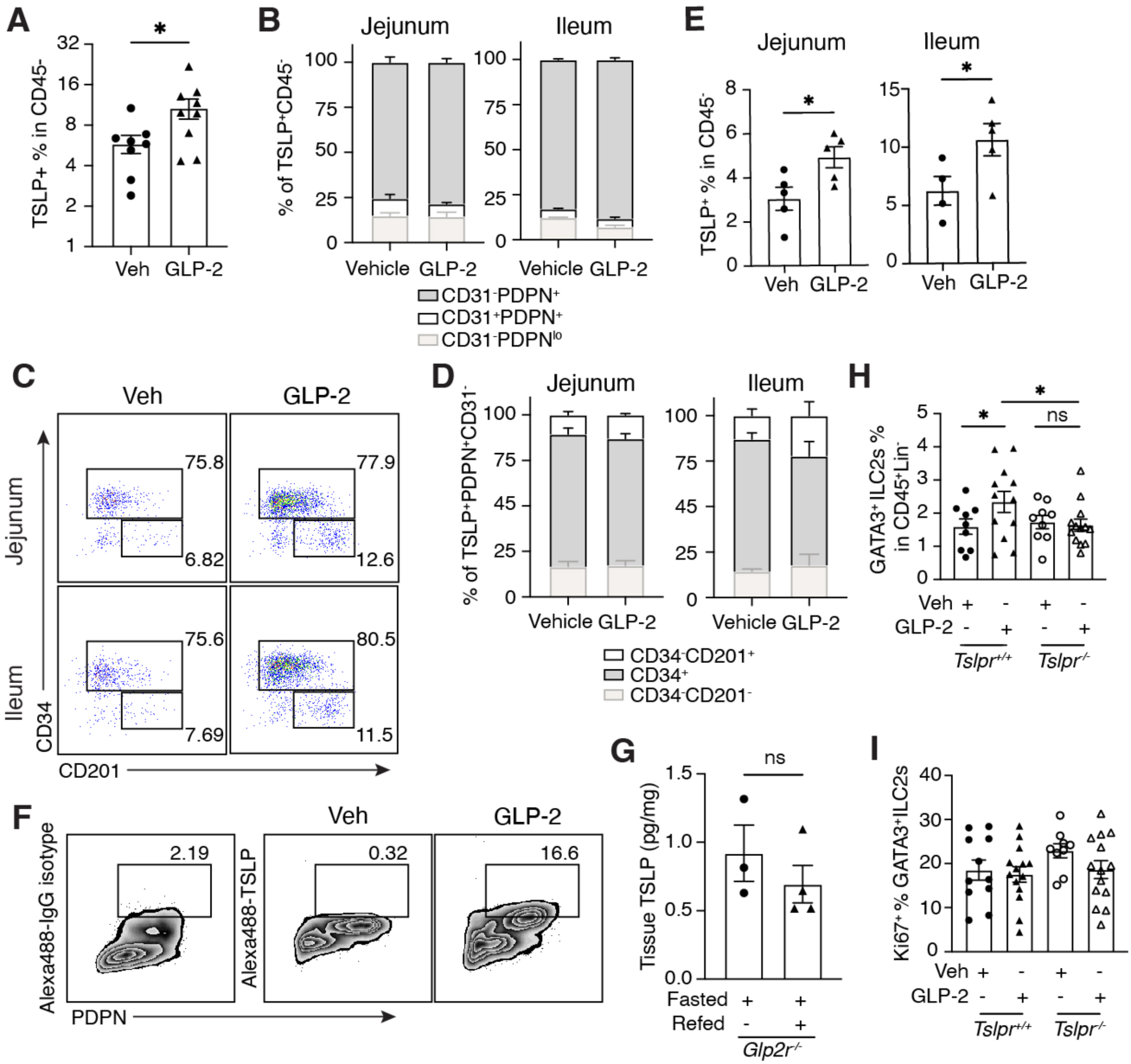
L cell GLP-2 drives GLP-2R- and TSLP-dependent ILC2 activation. (**A**)-(**D**) Flow cytometric analysis of TSLP-tdTomato^+^ cells in jejunal and ileal lamina propria (LP) at 2 hr after GLP-2[Gly2] injections. (**A**) Percentage of total TSLP-tdTomato^+^ cells among CD45^-^ cells in ileal LP. Veh, vehicle. (**B**) Proportion of endothelial and other stromal populations within TSLP-tdTomato^+^ cells in jejunum (**left**) and in ileum (**right**). **(C)(D**) Characterization of CD201^+^ and CD34^+^ cells. (**C**) Representative flow cytometric analysis. (**D**) Quantification of percentages of CD34^+^ and CD201^+^CD34^-^ within CD45^-^CD31^-^ TSLP-tdTomato^+^ stromal cells. (**E**) Quantitation of percentage of TSLP-tdTomato^+^ cells among CD45^-^ cells in jejunal (**left**) and ileal (**right**) LP after 3 daily injections of GLP-2[Gly2]. (**F**) Intracellular antibody staining for TSLP protein in purified CD45^-^CD31^-^TSLP-tdTomato^+^ stromal cells at 4 hr after incubation with 10 µg/ml GLP-2[Gly2] with Brefeldin A *in vitro*. (**G**) ELISA for TSLP protein recovered from ileal tissue homogenates from fasted *Glp2r^-/-^* mice after food gavage at 2hr (**H**)(**I**) Flow cytometric analysis of GATA3^+^ ILC2s in jejunal LP after 3 daily injections of GLP-2[Gly2]. (**H**) Quantification of percentage GATA3^+^ ILC2s among Lin^-^CD45^+^ cells. (**I**) Quantification of percentage of Ki67^+^ cells among GATA3^+^ ILC2s. ILC2s were gated on CD45^+^Lin^-^GATA3^+^ cells. All data are biological replicates, n≥3. Data are representative of at least two independent experiments. Error bars indicate samples mean± SEM. *P<0.03.

**Fig. S5.**
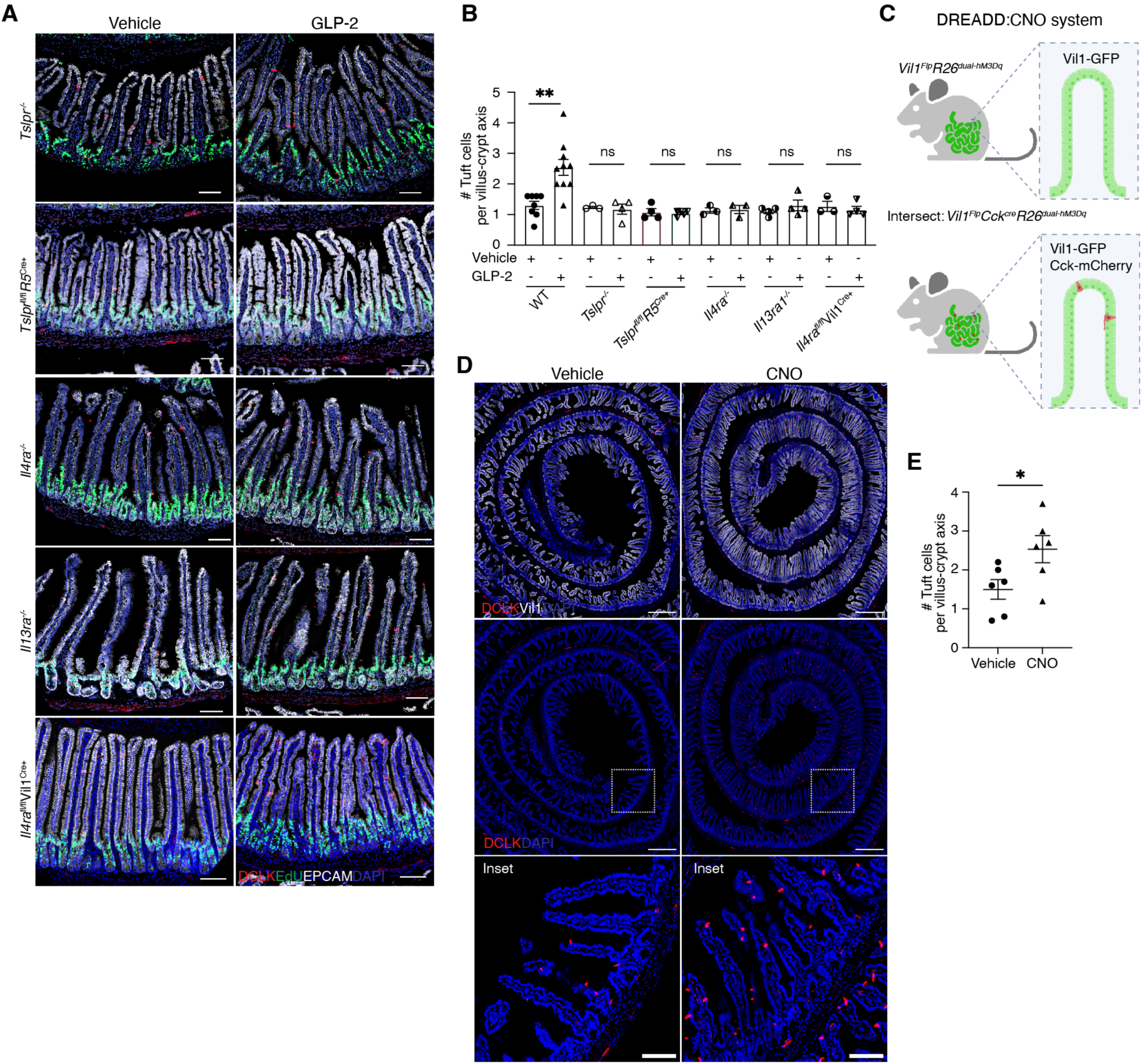
GLP-2 drives ILC2-dependent amplification of the tuft cell circuit. (**A**) Representative imaging of jejunal tuft cells in *Tslpr*^-/-^, *Tslpr^fl/fl^Il5^Cre+^ Il4ra*^-/-^, *Il13ra1*^-/-^ and *Il4ra^fl/fl^*Vil1^Cre^ mice. Green, EdU; Red, DCLK; White, EPCAM; Blue, DAPI. 20x objective. Scale bar, 100 µm. (b) Quantification of jejunal tuft cells after 3 daily GLP-2[Gly2] injections in WT, *Tslpr*^-/-^, *Tslpr^fl/fl^Il5^Cre+^, Il4ra*^-/-^, *Il13ra*^-/-^ and *Il4ra^fl/fl^*Vil1^Cre+^ mice. For tuft cell quantification, images were acquired using 20x objective and total DCLK^+^EPCAM^+^ cells in each field were counted and normalized by total number of villus-crypt axes in the field. Each dot represents a mouse. (**C**) Schematic illustrating the *Vil1^Flp^Cck^Cre^R26^Dual-hM3Dq^* DREADD and CNO system. (**D**) Representative imaging of jejunal tuft cells in *Vil1^Flp^Cck^Cre^R26^Dual-hM3Dq^* mice after administration of clozapine N-oxide (CNO) or vehicle control. Red, DCLK; White, Vil1-GFP; Blue, DAPI. 20x objective. Scale bar, 500 µm in the top and middle panels; 100 µm in the insets. (**E**) Quantification of jejunal tuft cells in *Vil1^Flp^Cck^Cre^R26^Dual-hM3Dq^* mice after administration of clozapine N-oxide (CNO) or vehicle control. Each dot represents a mouse.

**Movie S1-S7. Three - dimensional visualization of small intestine.**

Immunofluorescent staining of jejunum (Movie S1-S4) and ileum (Movie S5-S7) from Flare-TSLP mice. Red, TSLP-tdTomato; Green, PDGFRɑ; White, EPCAM; Blue, DAPI. 20x objective.

